# Epigenetic Intelligence: How Organisms Track Their Environment Through Molecular Memory

**DOI:** 10.64898/2026.01.08.698487

**Authors:** Holly V. Moeller, Hollie M. Putnam, Ross Cunning, Steven B. Roberts, Jose M. Eirin-Lopez, Roger M. Nisbet

## Abstract

The regulation of gene expression by epigenetic markers is a growing area of focus for researchers who seek to explain rapid phenotypic acclimatization to environmental change. Yet even as empirical datasets accumulate exponentially, the mechanistic underpinnings of epigenetic changes and their role in connecting environmental variation with the regulation of gene function remain unclear. Here, we use a stochastic model of epigenetic change to generate three testable predictions: (1) that organisms require environmental feedback to track their environments through epigenetic responses, (2) that this tracking requires coordination between the addition and removal of epigenetic markers, and (3) that epigenetically driven tracking is only effective under specific subsets of environmental variability. Despite the intuitiveness of these postulates, few to no experimental studies directly test them. We review correlational evidence consistent with the model’s predictions, describe hypothetical mechanisms by which epigenetic strategies could evolve, and clearly identify knowledge gaps and urgent experimental needs in the field. Overall, the field of environmental epigenetics is poised for major advances, which can be enhanced through the continued synthesis of mathematical frameworks and empirical data.

## 1 Introduction

Living organisms must maintain homeostasis in the face of variable environmental conditions. For example, long-lived animals may experience within-generational seasonal and interannual variability in temperature and resource availability, with direct impacts on metabolic function. One way in which organisms respond to both persistent and episodic environmental change is by shifting the sets of genes that they express to meet their contemporary needs (de Lamarck, 1809; Jablonka & Lamb, 1995). To maintain these shifts in expression patterns, many taxa employ “epigenetic modification,” in which modifiers are added to the genome to alter gene expression. These modifiers may include direct methylation of the DNA—including gene bodies (Sarda *et al*., 2012), and/or regulatory regions (Jones & Takai, 2001)—as well as modifications of the histone proteins that bind DNA and regulate its availability for expression (Marmorstein & Trievel, 2009) (Figure 1).

**Figure 1:**
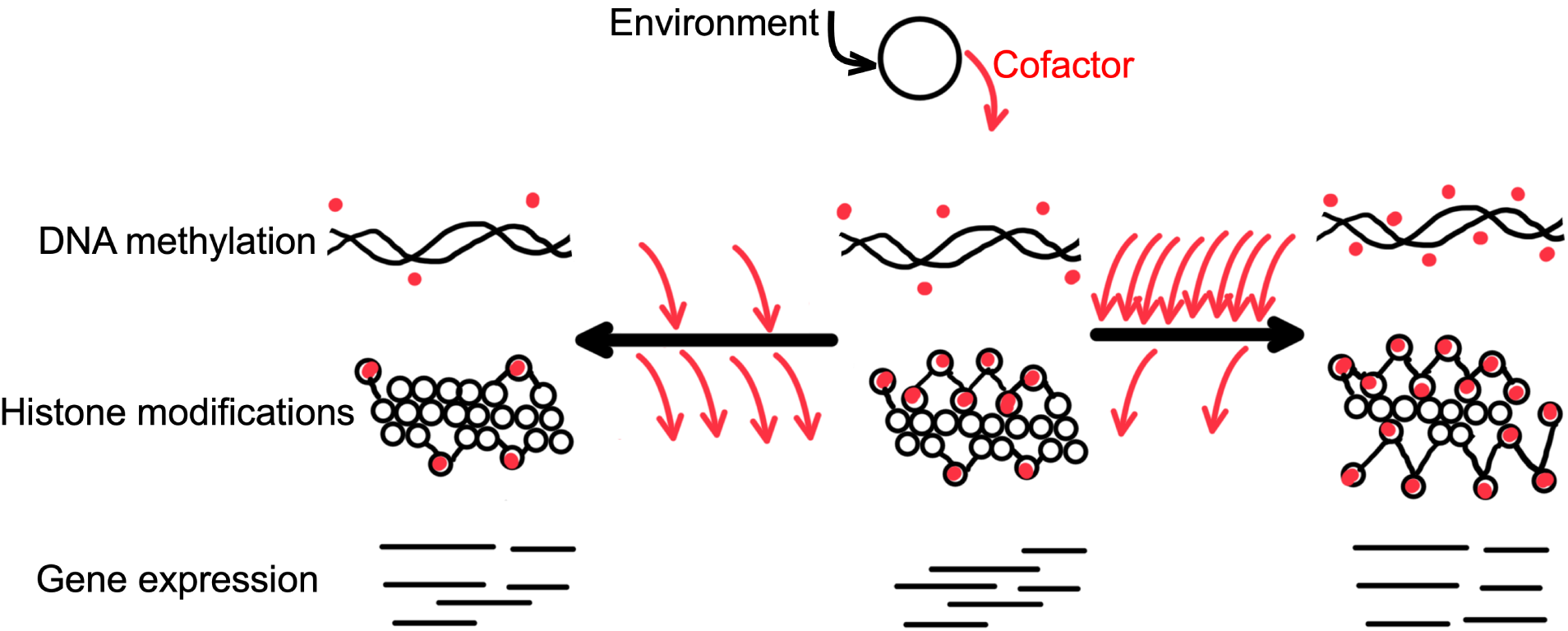
Mechanisms of epigenetic modification. When confronted with an environmental stimulus, organisms generate cofactors (red arrows) that can trigger epigenetic change. These cofactors act as substrates and/or catalysts of epigenetic modifications (red dots) including (1) DNA methylation (top row), in which methyl groups are added to cytosines, altering interactions between DNA, promoters, and RNA polymerases; and (2) histone post-translational modifications (middle row), in which DNA-binding proteins are modified in ways that alter their packing and, therefore, the exposure of DNA for transcription. The number of epigenetic markers placed in an organism’s genome depends upon the balance of both random and non-random addition and removal of these markers: when removal exceeds addition, there is a net decrease in markers (left-pointing black arrow and associated scenario), while relatively high amounts of marker addition produce the opposite (right-pointing black arrow). Although the effect of epigenetic modification on gene expression varies by lineage and genome location, the presence of these markers alters how DNA interacts with transcription factors and RNA polymerase, thereby shifting both the amount of gene expression and the splicing of exons (bottom row).

While the most detailed assessments linking epigenetic modification to phenotype have been carried out in traditional model systems (Jones & Takai, 2001), a growing body of evidence links epigenetics to the phenotypic plasticity of marine invertebrates (Eirin-Lopez & Putnam, 2019). Deepening this understanding is both urgent and critical: Marine invertebrates include many taxa important for human nutrition (e.g., oysters; Botta *et al*., 2020) and planetary biodiversity (e.g., stony corals; Eddy *et al*., 2021). They are also under increasing threat from human activity that is altering the thermal (Oliver *et al*., 2021) and chemical (Findlay *et al*., 2025; Kroeker *et al*., 2010) environments in which they are found.

Despite the immense promise of epigenetics as a mechanism to allow for rapid (within-generation) phenotypic adaptation to new environmental conditions, and a growing body of empirical data to be mined, few generalizable “rules” have emerged for how epigenetic modification may allow organisms to track their environment. Detailed mechanistic knowledge is still lacking about epigenetic modulation of gene expression in marine invertebrates (Baumgarten *et al*., 2018; Dixon *et al*., 2018; Li *et al*., 2018), particularly about the direct causal correlation between DNA methylation and differential gene expression (Abbott *et al*., 2024; Rodriguez-Casariego *et al*., 2022). Perhaps one of the most daunting tasks involves the integration of different types of information across biological levels (e.g., from epigenetics to gene expression to energetics to ecology), to make informed predictions and formulate hypotheses about the mechanistic basis underlying epigenetic responses and their implications for individuals and populations.

Mathematical frameworks can generate testable predictions about environmental triggers and mechanisms. Here, we develop a mathematical simulation that allows an organism to modify its epigenetic state to match (or fail to match) a changing environment. We use our mathematical model to draw broad conclusions about the information organisms need to track their environments, and the types of environments that can be epigenetically tracked. We link these model predictions with hypotheses about the mechanisms, function, and distribution of epigenetic environmental tracking, and briefly survey the existing literature to identify correlational (i.e., evidence consistent with our model’s hypotheses) and mechanistic (i.e., evidence supporting an explanatory linkage) information and knowledge gaps.

## 2 The Model

To identify the circumstances under which an organism can track its environment using epigenetic changes, we developed a simulation model that represents the iterative process of adding and removing epigenetic markers to an organism’s genome. These markers could represent changes in DNA methylation states, histone modifications, or other mechanisms leading to up- or down-regulation of gene expression (Eirin-Lopez & Putnam, 2019). Depending on environmental conditions, certain gene expression strategies may be more favorable than others (Barshis *et al*., 2013; Dayan *et al*., 2015); an organism that adjusts its epigenetic state to enable such favorable expression patterns will therefore be better matched to its environment (Jaenisch & Bird, 2003; Turner, 2009). Thus, organisms that are poorly matched to their environment should experience physiological benefits if they change their epigenetic markers to improve their match to the environment.

Here, we mimic this phenomenon by assuming that an organism experiences “stress” that is proportional to its phenotypic mismatch with local environmental conditions. The more stress an organism experiences, the greater the rate at which it remodels its epigenetic state. We first explore the epigenetic modification strategies that allow organisms to track their environments, and then mimic natural selection (by linking stress to fitness through the assumption that the organisms with the highest stress levels are most likely to die and that those with the lowest stress levels are most likely to reproduce) to find optimal (i.e., stress-minimizing) epigenetic modification strategies.

### 2.1 Model Components

#### Defining the Environmental and Epigenetic States

We represent the states of both the environment and the organism as E⃗, a binary string of “sites” that are either 1s or 0s (Figure 2). Each of the n positions in the string is a site that represents a hypothetical location in an organism’s genome that could take on one of two epigenetic state values in response to environmental conditions. The environmental string’s value at a particular position E*_i_* represents the organismal epigenetic state that would be best matched to the environment. A value of 1 indicates that an organism with a stress-minimizing epigenetic state *has* a marker in this location, and a 0 indicates that stress-minimization is achieved by the *absence* of an epigenetic marker.

**Figure 2:**
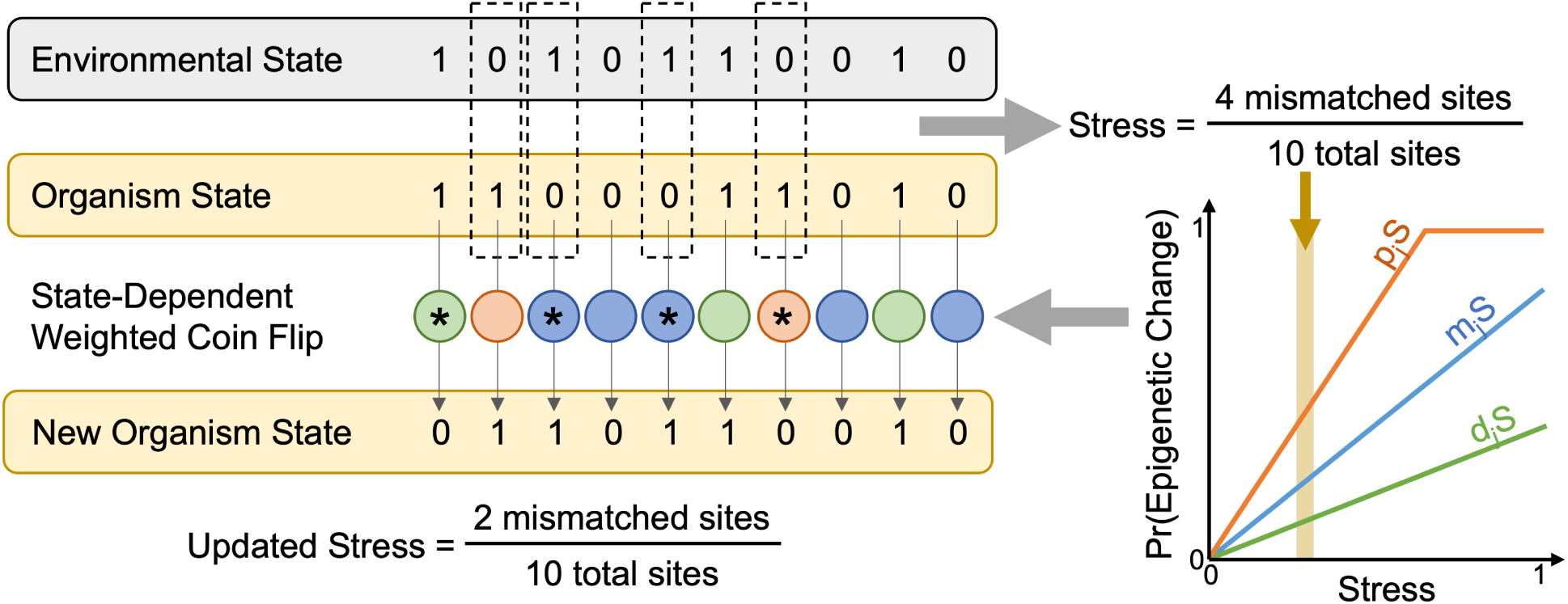
Conceptual diagram showing epigenetic modification. First, the organism’s state is compared to the environment and stress is computed. Based on stress levels and epigenetic change tendencies, the probability of an epigenetic change is computed. These probabilities are used to weight “coin flips” that determine which specific locations in the genome undergo random addition of epigenetic markers (blue), random removal of markers (green), or preferential removal of markers (orange). In the diagram, coin flips that result in epigenetic change are marked with a star. Following these changes, the organism’s stress can be recomputed (beginning the next timestep); in this case, stress has been reduced from 0.4 (4 mismatched sites) to 0.2.

Note that there are many possible interpretations of a site, which could represent, e.g., a specific cytosine which should be methylated (e.g., (Jones & Takai, 2001; Sarda *et al*., 2012)), or a gene or suite of genes whose expression is controlled by a cluster of epigenetic modifications made in concert (e.g., Adrian-Kalchhauser *et al*., 2020). Further, epigenetic markers may act to either repress or activate functional gene expression depending on organism, marker type, and gene regulatory network configuration. Therefore, our model merely postulates that marker presence or absence results in an environmental match, but is agnostic to the mechanism (e.g., activation or repression) by which that marker functions. Overall, we consider this string of epigenetic positions to represent the suite of an organism’s genes that can be environmentally responsive; That is, we are neglecting genes that are silent or constitutively expressed, or that have invariant epigenetic states.

The environment can vary. There are various ways of implementing environmental change (i.e., shifts of E*_i_* from 0 to 1 or vice versa). We consider two possibilities: In one case, we periodically redraw the entire environmental string, which represents an “extreme event” with an abrupt and random perturbation. There is therefore no correlation between the past epigenetic optimum and the current one following the perturbation. In the second case, we allow the environment to vary cyclically by gradually shifting the environmental epigenetic state over time in a systematic manner. In the latter case, we can control both the speed (frequency of shifts) and magnitude (number of changed sites) of environmental change, allowing us to mimic cyclic perturbations like seasonality, interannual variation, and decadal cycles.

#### The Organisms

We use an individual based model to represent the population of living organisms in our model. Each individual is represented by its own binary string, O⃗*_j_*, of the same length as the environmental string. Each site on an organism’s string corresponds to the same site in the environmental string; this allows an individual organism to experience variable degrees of “stress” depending on the number of sites that are mismatched with the environment. Our model posits that an organism’s epigenetic state can undergo directional change gradually, through the accumulation of a few directional (i.e., information-based) marker removals amidst a largely random set of additions and removals. In this way, our model mimics the effort of cells to harness chemical entropy to generate directed chemical forces (Baker, 2023). An increasing intensity and duration of environmental signals will promote increasing concentrations of cofactors available for the epigenetic regulation of genome function as a response. We describe a mechanistic representation of this process below.

#### Stress-Dependent Epigenetic Change

We assume that organisms undergo epigenetic modification when confronted with detectable physiological stress (Hackerott *et al*., 2021), which we here assume stems from a mismatch with their environment. We define stress as the average degree of mismatch with the environment across all n possible epigenetic sites. Individual j’s level of stress S*_j_* is therefore given by:

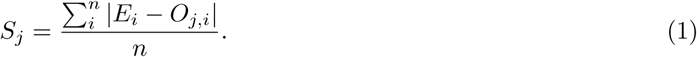

Note that this approach assigns each epigenetic site equal weighting, and assumes that incorrectly marked sites (i.e., organism = 1, environment = 0) and incorrectly unmarked sites (i.e., organism = 0, environment = 1) are equivalently stressful.

We assume that an organism makes epigenetic modifications proportional to the level of stress it is experiencing. In other words, if an organism is well-matched to its environment, its stress levels are low, and it will make few, if any, epigenetic changes. But when an organism’s stress levels are high, it will increase rates of epigenetic remodeling in an attempt to more quickly achieve environmental optimality. Therefore, we represent each organism’s probability of adding an epigenetic marker to a single site (i.e., changing that site’s epigenetic state from a 0 to a 1, such as via methylation) during one time step as proportional to its current stress level S*_j_*, scaled by an individual-specific marker addition tendency m*_j_*:

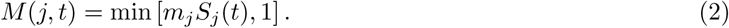

This stress-dependent probability is used to determine whether or not a currently unmarked site is marked during a given round of epigenetic change. Specifically, for each unmarked site, the algorithm draws a value between 0 and 1 from a uniform distribution. If this value is greater than the marking probability M(j, t), an epigenetic marker is added. Per-round probabilities are capped at 1. Statistically, this means that the total number of sites that are marked in a given round of epigenetic modification is drawn from a binomial distribution with a probability M(j, t) and a number of trials equal to the number of presently unmarked sites (state = 0).

The removal of markers is scaled similarly, with an individual marker deletion tendency d*_j_*:

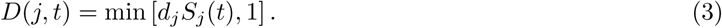

3

Addition and deletion probabilities tell us how likely an organism is to remodel each position in its string O⃗*_j_* at each timestep. However, these probabilities may change if the organism has information about which sites are currently mismatched with its environment. Such environmental information might be collected when, for example, excess gene products inhibit gene expression. We use a scalar to increase the probability of “correct” epigenetic changes. Thus, the probability of *preferentially* deleting an epigenetic marker from a mismatched site (i.e., a location where the environmental state is 0 and the organismal state is 1) is given by:

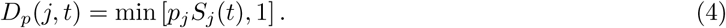

where p*_j_* is the individual-specific preferential removal tendency. Although both addition and removal can occur preferentially, here we consider only preferential removal. However, our results are robust to the alternate assumption (Figure S1).

Because similar pools of cofactors and cellular equipment are likely used for both preferential and random deletion of epigenetic markers, we assume that these two probabilities must be related to one another. We therefore assume that there is an overall marker removal tendency o*_j_*, that is governed by the production of such machinery, and that an environmental information parameter, ɛ, links the random and preferential change tendencies, which are centered around this overall tendency:

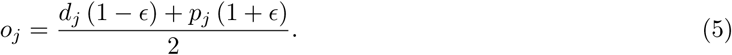

Here, we choose 0 ≤ ɛ ≤ 1 so that all removal probabilities are nonnegative.

It is convenient to link the addition and removal of epigenetic markers, which we do with the parameter b, which scales the tendency of removal relative to the tendency of addition. Because b = m*_j_*/o*_j_*:

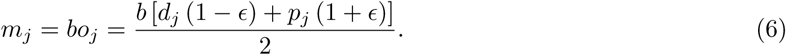

If b > 1, marker removal is more stress-sensitive than marker addition, and if b < 1, the opposite is true.

### 2.2 Algorithm Steps

Simulations proceed through a series of rounds. At the start of each round, for each individual, we (1) compute the individual’s stress, (2) probabilistically add epigenetic markers, and (3) probabilistically remove epigenetic markers (Figure 2). Following these individual epigenetic updates, any environmental change takes place via an adjustment to the E⃗ vector. Because epigenetic updating occurs at every round, and more frequently than birth-death events (when included; see below) or large-scale environmental change, our model best represents within-generation epigenetic modification timescales.

### 2.3 Model Analysis

We used our model to address three questions about how epigenetic modifications could allow organisms to track their environment. We asked (1) whether organisms must receive some information about the state of the external environment in order to be able to epigenetically track the environment; (2) what relative rates of addition and removal of epigenetic markers allow an organism to minimize its stress by tracking its environment; and (3) under which environmental conditions epigenetic change is an effective mechanism for environmental tracking. Below, we describe our analytical approach and the model’s predictions for each query. We compare the model’s predictions with existing empirical knowledge, and identify key research priorities to address knowledge gaps.

## 3 Results

### 3.1 Tracking the environment requires site-level information

How do organisms gather information about the environment around them, and is this information necessary for them to tune their epigenetic state to match environmental needs? While stress from environmental mismatch can be measured at the aggregate scale, it is less clear how organisms might identify which specific epigenetic changes should be made based on environmental cues. Therefore, we first used our model to investigate what information organisms need to detect about their (mis)match to the environment in order to track the environment using epigenetic mechanisms.

#### 3.1.1 Model Results: The mathematical model predicts information-dependent tracking

We examined the ability of an organism to track a stochastically changing environment as a function of its degree of information about the state of that environment. Specifically, we varied the magnitude of the difference between preferential and random marker removal probabilities as a proxy for the amount of information an organism has about its environment. When no information is present, all changes to epigenetic state occur randomly, such that ɛ = 0 and d*_j_* = p*_j_* = m*_j_*/b*_j_*. When all marker removal is information-based, the only sites chosen for removal are those that produce matches to the environment. In this case, ɛ = 1, such that d*_j_* = 0 and p*_j_*= 2m*_j_*/b*_j_*.

We found that only organisms that can obtain environmental information are able to track the environment (Figure 3). These organisms experience transient increases in stress when the environment changes, triggering a period of marker removal (Figure 3b). Over time following the environmental perturbation, these organisms adjust their epigenetic state to match their environment, returning their mean number of epigenetic markers to the pre-disturbance state and lowering their stress levels from ∼ 0.5 (the mean expected proportion of mismatched sites when the environment and organism are drawn randomly) to values as low as 0.15 (a 15% mismatch with their environment).

**Figure 3:**
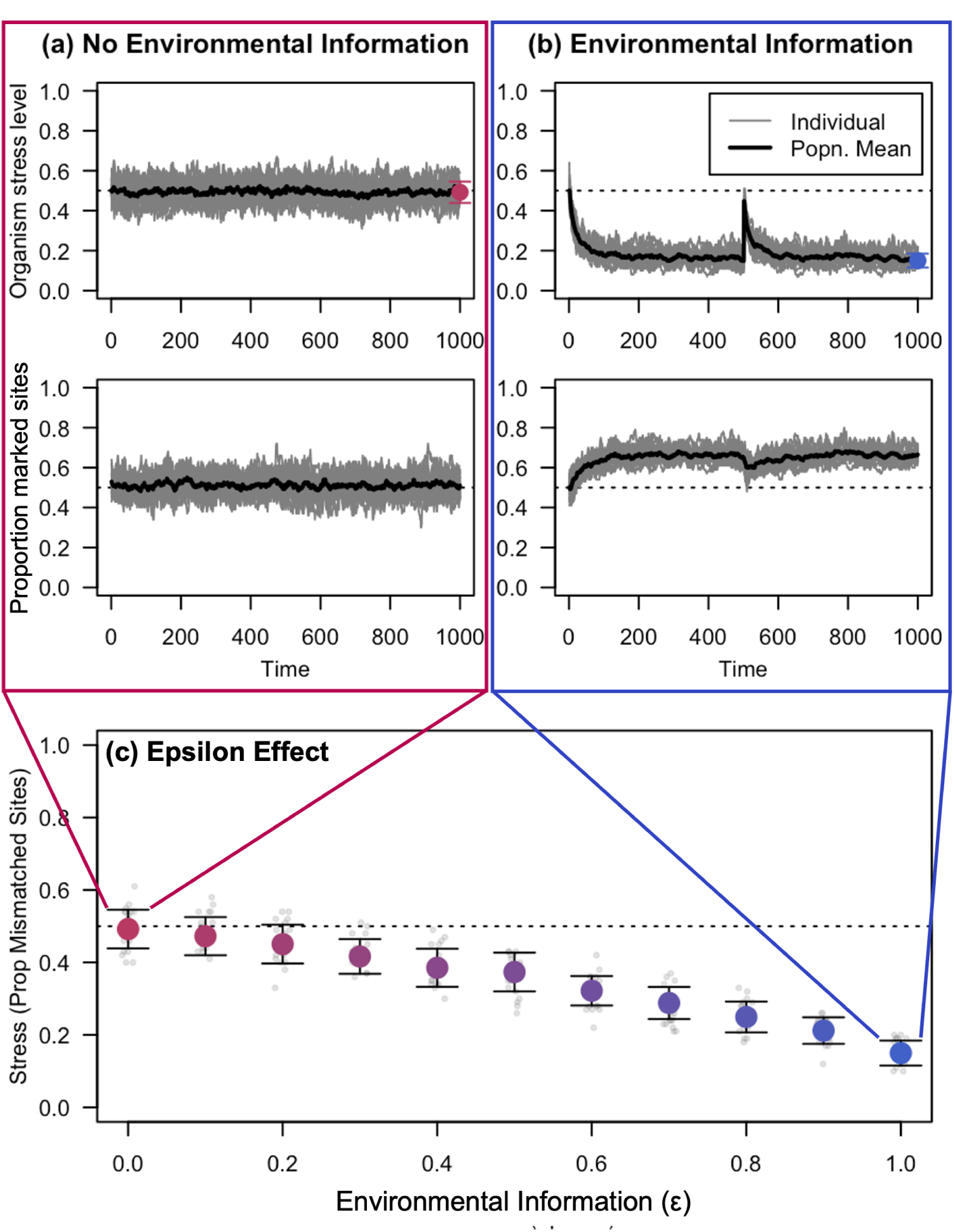
Effects of environmental information on organismal stress. Top: Timeseries showing a group of 20 organisms tracking (or failing to track) their environment over time. Without environmental information, stress levels hover around the mean expectation of 0.5 (a), but with environmental information, the organisms are able to reduce their stress levels within about 100 timepoints (b). Bottom: The larger the difference in baseline and preferential marker removal tendencies, the better the organism’s performance. Simulations are for 20 organisms with a baseline marker addition tendency of 0.2 tracking an environment with a mean epigenetic marker abundance of 0.5. Gray points show individual stress levels, and colored points and error bars show population means and standard deviation.

The magnitude of stress reduction depends on how successfully organisms can decouple random (d*_j_*) and preferential (p*_j_*) marker removal (Figure 3c). The optimal solution would be to set d*_j_*= 0 so that no random marker removal occurs, and then to increase p*_j_* without bound so that the probability of accurately removing markers from mismatched sites is high, even when stress is low. However, it seems unlikely that an organism would be able to completely avoid inadvertent marker removal: First, random marker removal is likely during cell division because epigenetic markers are not always copied over to new strands of DNA (Edwards *et al*., 2017). Second, when there is high production of epigenetic cofactors (e.g., modification enzymes) related to stress response (i.e., if p*_j_* is large), it seems reasonable to suppose that some inadvertent marker removal will also take place. For example: if an organism has produced a lot of cellular enzymes that demethylate cytosines, then these enzymes will likely increase both preferential (i.e., environment-informed; p*_j_*) and random (d*_j_*) marker removal tendencies. Thus, in our subsequent analyses, we coupled these rates to each other by setting ɛ = 0.8.

#### 3.1.2 Correlational Evidence: Empirical studies demonstrate epigenetic responses to environmental change

What empirical evidence exists that marine invertebrates use environmental information to directionally shift their epigenetic states in order to track the environment? Numerous studies have shown that organismal phenotypes may vary predictably across environmental gradients (Morel-Journel *et al*., 2020), and may shift dynamically in response to seasonal changes (Shearer *et al*., 2016; Suarez-Ulloa *et al*., 2019), and directionally in response to perturbations like thermal and irradiance stress (Kelly *et al*., 2017) (Gomez-Campo *et al*., 2024).

In marine invertebrates, a growing body of evidence is beginning to link these phenotypic changes to epigenetic changes by examining how epigenetic state varies in response to acute environmental shifts and fluctuating environmental conditions. For example, adult oysters from the genus *Crassostrea* respond to ocean acidification (OA) through epigenetic changes in mantle tissue (Chandra Rajan *et al*., 2021; Downey-Wall *et al*., 2020), reproductive tissue (Venkataraman *et al*., 2020, 2022), and larvae (Dang *et al*., 2023; Lim *et al*., 2021). In the Pacific Oyster, infection with Pacific Oyster Mortality Syndrome resulted in differential DNA methylation linked to susceptibility and resistance to the syndrome (Valdivieso Munoz *et al*., 2024). In geoduck clams exposed to OA, changes in DNA methylation are correlated with changes in shell size, with both methylation and phenotypic changes persisting for months even after the environmental perturbation ends (Putnam *et al*., 2022). Further global DNA methylation changes have been documented in the coral *Pocillopora acuta* following exposure to low pH for 6 weeks (Putnam *et al*., 2016).

Some longer-term studies in marine invertebrates have also demonstrated variability in epigenetic state linked to phenotype. For example, studies in the flat tree oyster *Isognomon alatus* revealed seasonal changes in DNA methylation patterns over a 2-year period, which were primarily associated with changes in temperature and horizontal visibility (Suarez-Ulloa *et al*., 2019). Studies have also demonstrated hypomethylation following acute exposure to low pH in the Antarctic pteropod, *Limacina helicina antarctica*, which then returned to normal levels over time (Bogan *et al*., 2020), consistent with our model’s prediction of transient loss of epigenetic markers following an environmental perturbation (Figure 3C).

#### 3.1.3 Mechanistic Evidence: What mechanisms might allow organisms to translate environmental information into epigenetic modifications?

Consistent changes in epigenotype and phenotype with environmental change suggest a mechanistic linkage between an organism’s environment and its epigenetic state (e.g., that some epigenetic modifications occur non-randomly, as in our model). Yet the mechanisms by which organisms use environmental cues to modify gene expression networks remain unclear. Some mechanistic linkages have been shown in vertebrates. For example, in the European sea bass, methylation patterns of the cyp19a promoter correlate with temperature shifts, suggesting that epigenetic modifications may mediate the effects of temperature on sex differentiation (Navarro-Martín *et al*., 2011; Piferrer, 2013)

In corals, epigenetic responses to ocean acidification may involve epigenetic modifications in pathways controlling skeletal calcification processes (Liew *et al*., 2018). Exposure of the coral *Stylophora pistillata* to reduced pH (7.2, 7.4, and 7.8 relative to 8.0) for ∼ 2 years resulted in differentially methylated genes in the mitogen-activated protein kinase signaling pathway (specifically, MAPK signalling regulators MAPK and JNK), driving changes in cell size and shape that led to differences in coral skeleton growth and porosity (Liew *et al*., 2018). Further, in the staghorn coral *Acropora palmata*, changing light conditions using reciprocal transplantation for 13+ weeks resulted in photoacclimation and skeletal change linked to differences in gene body DNA methylation (Gomez-Campo et al 2023). In the model marine cnidarian *Aiptaisa*, epigenetic modifications of DNA methylation act to reduce spurious transcription; these are established and maintained by active transcription triggering histone modifications (trimethylation of histone H3 at lysine 36; H3K36me3) (Li *et al*., 2018). This histone modification includes a mark that binds to a specific protein domain in the *de novo* DNA methyltransferase, Dnmt3b, which in turn supports ongoing DNA methylation. Disentangling the cascade of epigenetic events requires experiments that concurrently quantify physiology, gene expression and epigenetic mechanisms at multiple time points, rather than an endpoint approach.

Understanding the generality of epigenetic mechanisms across taxa and stressors requires first determining how different epigenetic markers (in different regions of the genome) act to alter gene expression. For example, while in vertebrates methylation is generally understood to act as an on/off switch for gene expression (Aquino *et al*., 2018), in invertebrates the directionality is less clear (Roberts & Gavery, 2012). Additionally, while the links between gene body methylation and the magnitude and/or variation of gene expression are consistent across studies (Eirin-Lopez & Putnam, 2019; Kim *et al*., 2024), the mechanisms underlying these correlations are unexplored. Further, hypotheses linking DNA methylation to alternative splicing (Roberts & Gavery, 2012) remain underexplored in marine invertebrates, as well as the cross-talk of epigenetic multiple mechanisms (Adrian-Kalchhauser *et al*., 2020). Closing these knowledge gaps will require additional mechanistic studies linking environmental change to epigenetic state (Figure 4). Ideally, these studies should identify the types and timing of epigenetic changes triggered by known environmental changes, as well as the cofactors that mediate these changes. Collectively, identifying putative mechanistic pathways will allow us to understand the full scope of potential epigenetic alterations and, in turn, the magnitude and speed of environmental change that organisms may be able to withstand.

**Figure 4:**
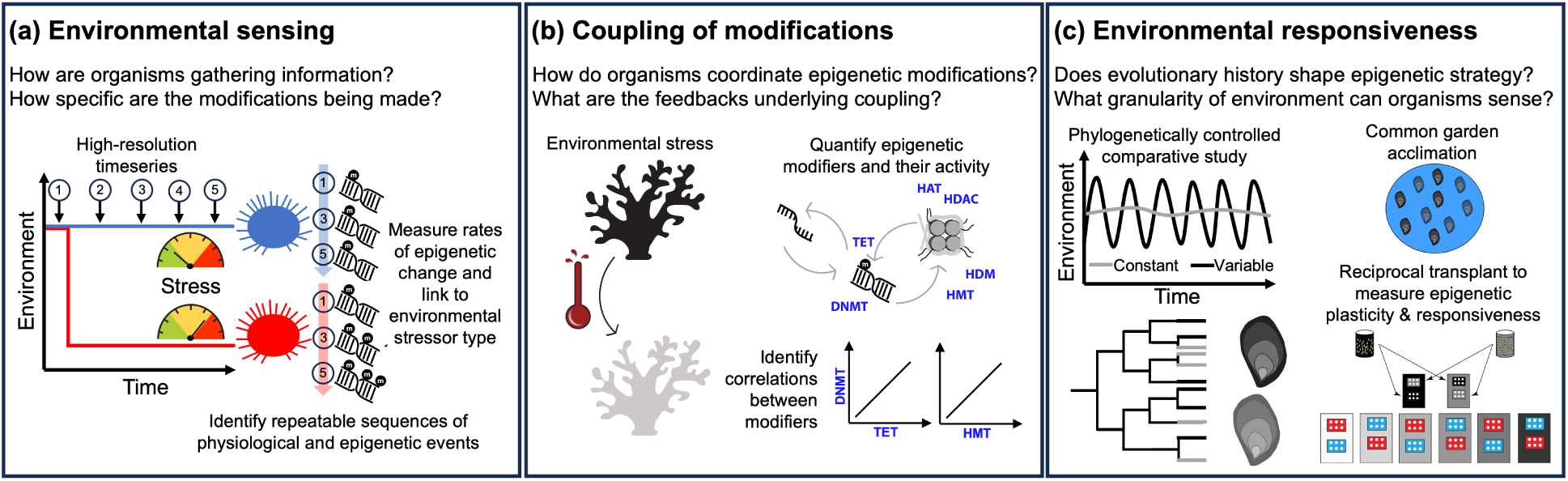
New experimental studies are necessary to address knowledge gaps. Here, we propose three categories of experiments: (a) To identify the mechanisms by which marine invertebrates use environmental information to modulate epigenetic change, high-resolution timeseries data can be collected following an experimental manipulation to induce epigenetic change. Ideally, experiments would be repeated with multiple lineages and different species to identify any common mechanisms. (b) To determine how organisms maintain balanced amounts of epigenetic modifications, similar experiments could quantify the amount and activity of enzymes (blue text labels) that catalyze epigenetic modifications following stress. (c) Finally, to test the model’s predictions about environments that should select for epigenetic modification strategies, comparative approaches could explore the epigenetic strategies of lineages that have evolved in comparatively stable or variable environments. By first acclimatizing lineages to a common garden, and then exposing them to environmental stressors with varied periodicity, both the *magnitude* of the epigenetic response and its *sensitivity to the frequency of environmental change* can be estimated.

### 3.2 “Optimal” coupling of epigenetic change tendencies depends on the environment

The dynamic process of adding and removing epigenetic markers allows for the location of these markers to shift over time, enabling an organism to respond to changes in its environment. However, in order to maintain an equilibrium mean level of epigenetic markers comparable to its environment, an organism’s loss and addition of these markers must be roughly balanced. Therefore, we next studied how marker addition and removal tendencies (which in our model underlie the linkage between an organism’s stress and its rate of epigenetic change) must be coupled to one another in order to allow for epigenetic tracking of the environment.

#### 3.2.1 Model Results

If marker addition and removal probabilities become too decoupled, an organism could lose all epigenetic markers (if b >> 1 such that d*_j_*, p*_j_* >> m*_j_*) or add markers to all of its genome (if b << 1 such that m*_j_* >> d*_j_*, p*_j_*). In an environment with an average marker abundance of 0.5, an organism that has lost (or gained) all of its epigenetic markers experiences a high level of stress (0.5). Therefore, the more mismatched the marker alteration tendencies, the higher the stress level (Figure S2).

We computed optimal pairs of marker addition and removal tendencies by fixing the former and using a pseudo-evolutionary algorithm to efficiently find values of the latter that minimized organismal stress. Specifically, we modified our algorithm to include birth-death processes that eliminated the highest stress individual and replaced that individual with a duplicate of the lowest stress individual (mimicking reproduction of the highest fitness individual). For maximum simplicity, we assume that birth events duplicate the parent organism’s parameters (e.g., m*_j_* and d*_j_*), as well as its present epigenetic state O⃗*_j_*. Birth-death processes occurred every 10 rounds of epigenetic change, and every tenth birth-death event also involved a mutation in the marker removal tendency. When a mutation occurred, a new mean marker removal tendency was drawn from a normal distribution with standard deviation 0.1 centered around the parent’s removal tendency. We bounded the distribution at zero, so that mutations that resulted in m*_j_* < 0 or d*_j_*< 0 were rounded to m*_j_* = 0 or d*_j_* = 0, respectively. Thus, over time, populations evolved marker removal tendencies that minimized stress (Figure S3).

Our results highlight the importance of coupling epigenetic marker regulatory tendencies. Generally, higher marker addition tendencies necessitate higher marker removal tendencies so that the average level of genome marking remains constant (Figure 5). Decoupling these rates produces high stress either because of rampant marker removal (if o*_j_*> m*_j_*) or addition (if o*_j_* < m*_j_*) (Figure S4). However, the exact coupling of these tendencies depends upon environmental conditions. For example, when the environment requires relatively few epigenetic markers, marker removal tendencies must be relatively greater in order to keep the organism’s overall amount of epigenetic markings low (Figure 5a), but when the environment requires a high amount of epigenetic markers, organisms should evolve relatively low removal tendencies (Figure 5).

**Figure 5:**
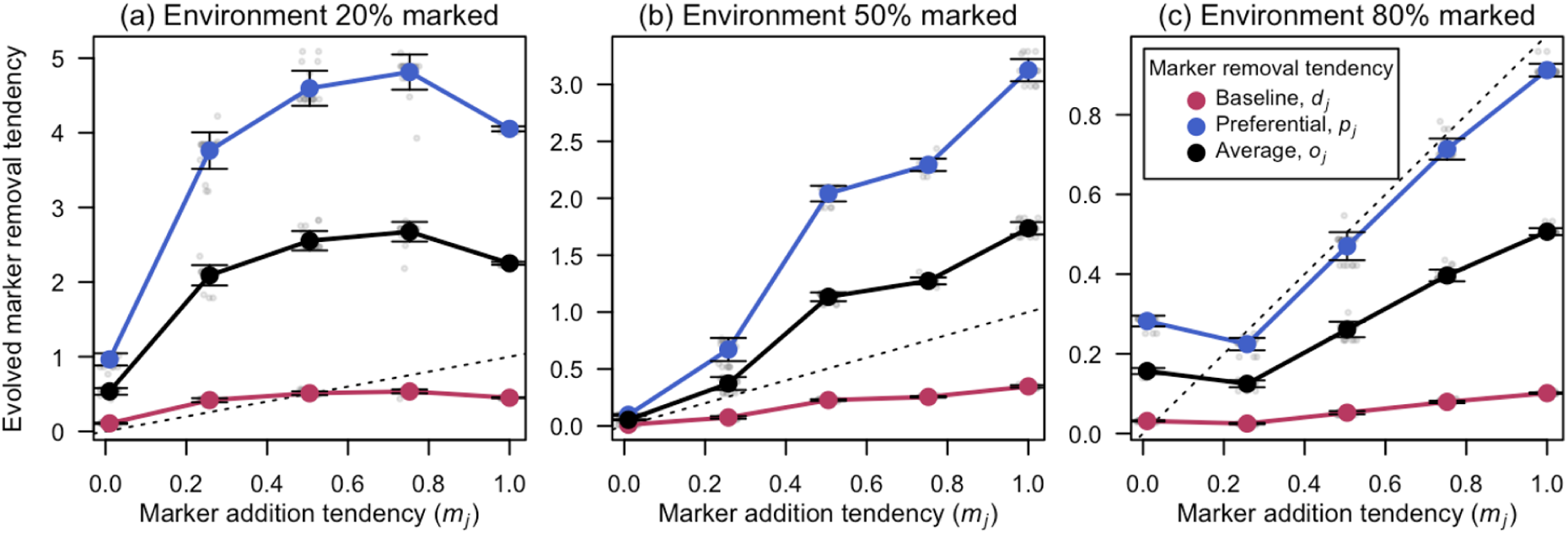
Evolved stress-minimizing marker removal tendencies as a function of (fixed) addition tendencies. (Note that baseline and preferential marker removal are coupled by ɛ = 0.8.) Generally, as an organism’s tendency to add markers in response to stress increases, its tendency to remove markers must increase proportionately. The more markers in the environment that the organism is tracking, the lower the evolved marker removal rates (compare from left to right, noting differences in y-axis scales).

#### 3.2.2 Correlational Evidence: Organisms appear to maintain homeostasis in epigenetic markers over long timescales

Consistent with our model’s prediction that organisms should re-equilibrate at a consistent level of epigenetic markers following environmental change (Figure 3AB, Figure S2), data from the Arctic Pteropod, *Limacina helicina antarctica* shows transient reductions in DNA methylation in response to one and three days of acute low pH exposure, and then recovers to the same levels as ambient by 6 days of exposure (Bogan *et al*., 2020). Similarly, in the reef-building coral *Acropora cervicornis*, DNA methylation profiles diverged following 2 days of exposure to increased temperature in four different heat accumulation treatments (2, 4, 5, and 9 Degree Heating Days) compared to corals at ambient temperature (29°C), but were indistinguishable when measured eight days after the heat stress terminated (Martell *et al*., 2024).

#### 3.2.3 Mechanistic Knowledge Gap: What mechanisms ensure coupling of addition and removal of epigenetic markers?

In order to return to and maintain a consistent level of epigenetic markers, organisms must balance the addition and removal of these markers. Our model proposes a stress-mediated feedback based on environment-performance mismatch that organisms could use to coordinate epigenetic modifications, but the exact mechanism by which this coordination takes place remains unclear. It is known that the addition and removal of markers are catalyzed by distinct enzymes: for example, histone acetyltransferase (HAT) and deacetylase (HDAC) regulate the addition of histone modification tags, while lysine- and arginine-specific histone demethylases (HDM) regulate their removal (Marmorstein & Trievel, 2009). Further, a series of DNA methyltransferases (DNMT) and ten-eleven translocation (TET) enzymes regulate the addition and removal of methyl groups from cytosines (Li & Zhang, 2014). Therefore, organisms are able to shift their overall levels of epigenetic markers by altering the relative ratios of cofactor pools.

Epigenetic modification is an energy-requiring process, both for the production of the enzymes necessary to catalyze these reactions and for the high-energy substrates that are needed for the addition and removal of some markers (Donohoe & Bultman, 2012). Thus, one way that epigenetic modification may be linked to environmental stress is through an organism’s energetic state (e.g., Berger *et al*., 2025). Organisms may only be willing to invest energy in modifying their epigenetic state if they perceive a costly (in terms of energy gain and/or fitness) mismatch with their environment. This also implies (we hypothesize) that organisms that are in a higher energy state (e.g., due to stochastic processes related to resource acquisition, or because they are better able to track their environment quickly) may be best able to track a changing environment. To understand how rates of epigenetic marker addition and removal are coupled to one another, future studies should examine the quantities of enzymes and substrates, and their activity, over time during an environmental change event and its aftermath (Figure 4B).

### 3.3 Tracking is only effective in certain environments

Although epigenetic modifications can, in principle, allow organisms to become locally acclimatized to their environment, we do not know the range of environments that can be effectively tracked via epigenetic mechanisms. Therefore, we next sought to identify how organismal stress levels varied with different types of environmental change and, therefore, what stress-minimizing epigenetic modification strategies might maximize performance.

#### 3.3.1 Model Results

We used our model to explore the relationship between the frequency and magnitude of environmental change and organism stress. To do this, we first implemented a new algorithm for environmental change that allowed the environment to change with predictable fluctuations. Specifically, we simulated a “band” of optimally methylated sites that shifted in genome location with a fixed frequency (per number of simulation timesteps) and magnitude (number of genome sites). Examples of how this model can be tuned from low-frequency, large changes to high-frequency, small changes can be seen in Figure 6. We used results from our evolutionary simulations (Figure 5b, for a 50% marked environment) to optimally couple marker addition and removal tendencies. Specifically, we simulated a population of individuals with a marker addition tendency m*_j_* = 0.1 and a mean marker removal tendency o*_j_* = 0.21.

**Figure 6:**
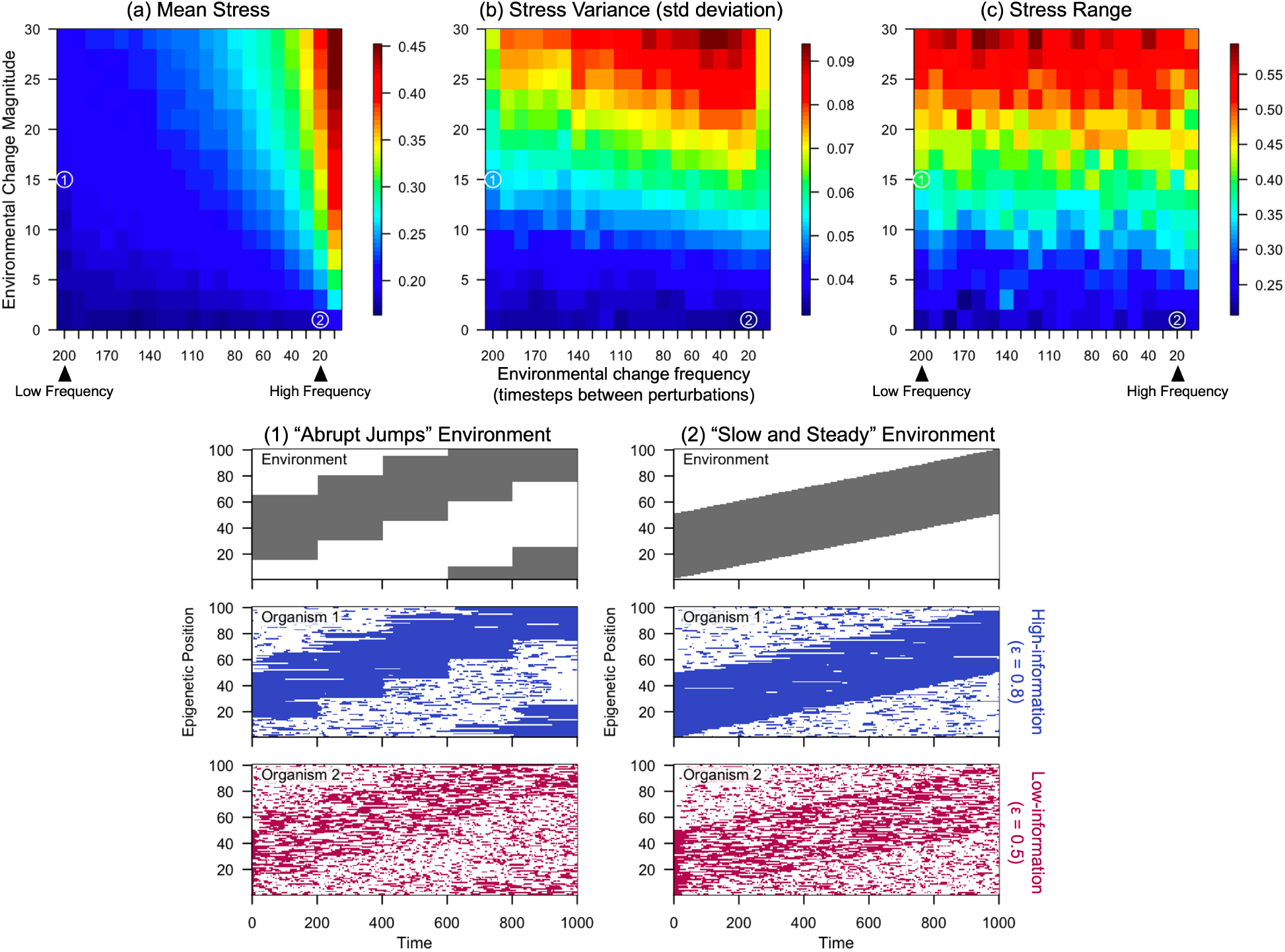
Effects of environmental change on organism performance. (a) An organism with strong environmental tracking (ɛ = 0.8; m*_j_* = 0.1 and o*_j_* = 0.21) is able to track environments that change with low frequency or low magnitude. (b) However, high-magnitude changes result in large variance in stress levels, because of transient periods of re-equilibration to environmental conditions immediately following a shift in the environmental state. When changes are frequent and high-variance, stress variance actually decreases because of the extremely poor match to the environment. (c) Overall, larger magnitude shifts produce greater range in stress (comparing minimum to maximum values over the course of the simulation) because of periodic spikes in stress level. Below: Example scenarios (also marked on the heatmaps) for (1) low-frequency, intermediate-amplitude, and (2) high-frequency, low-amplitude environmental change. Filled area indicates a site with an epigenetic marker in the environment (top row, gray), an organism with a high degree of tracking (middle row, blue), and an organism with a low degree of tracking (bottom row, pink). Organisms with high levels of environmental information (ɛ = 0.8) track better, especially in environments with larger magnitudes of change.

When environments change relatively infrequently (e.g., once every 200 timesteps; Scenario 1 in Figure 6), organisms with high-information tracking strategies (e.g., ɛ = 0.8) are able to shift their epigenetic state alongside environmental change, minimizing stress even if the environmental change is large in magnitude. However, frequent environmental changes can only be tracked if they are small in magnitude (e.g., Scenario 2 and upper right corner of Figure 6), and otherwise produce high levels of stress that are comparable to a random epigenetic strategy (expected mean stress of 0.5).

Our model does not consider any costs for epigenetic modifications, so an organism that attempts to track its environment cannot incur any fitness costs in excess of its mismatch. Thus, our model cannot be used to demonstrate tradeoffs between epigenetic tracking and energy conservation, and does not have selection pressure to reduce epigenetic modification. Incorporating such costs could, however, reveal “optimal” epigenetic modification strategies, including abandoning epigenetic change when environmental tracking benefits are less than energetic costs.

The faster the environment shifts (either via high frequency changes or large magnitude changes), the more (accurate) epigenetic changes must be made to track it. Thus, only organisms with sufficiently high tendencies to preferentially add or remove epigenetic markers are capable of tracking the environment (Figure S5).

#### 3.3.2 Correlational Knowledge Gap: Do organisms from more variable environments rely more on epigenetic modifications?

Our model leads to the hypothesis that epigenetic regulation of gene expression is a more effective tool in some environments than others (Figure 6). For instance, organisms in highly variable environments (e.g., intertidal zones, estuaries, seasonal environments) often face fluctuating temperature, salinity, oxygen levels, and food availability, with epigenetic mechanisms likely playing a role in necessary phenotypic plasticity. Epigenetic modifications can provide a flexible and rapid way to adjust physiology through gene expression. Indeed, there is strong evidence that highly variable environments impact marine invertebrate phenotype (Hsieh *et al*., 2024; Keppel *et al*., 2016; Pernet *et al*., 2007; Safaie *et al*., 2018) and direct their gene expression profiles, including expression plasticity (Kenkel & Matz, 2016) and front-loading (Barshis *et al*., 2013; Brown *et al*., 2025; Collins *et al*., 2021; Gurr *et al*., 2022).

Direct tests of the hypothesis that epigenetic regulation of gene expression is a more effective tool in some environments than others are challenging because organisms from different environments are typically also separated by evolutionary history with potential stochastic impacts on epigenetic strategies. However, comparative studies, in which closely related organisms from environments with different degrees of variability, could be used to test this hypothesis. For example, oysters (genus *Crassostrea*) provide a good system due to their broad environmental distributions and tolerance to highly variable conditions. Species such as the Eastern oyster (*C. virginica*) and the Pacific oyster (*C. gigas*) inhabit estuarine environments that experience frequent fluctuations in temperature, salinity, and oxygen availability, whereas other closely related species (e.g., *C. nippona*) occupy more stable coastal environments (Hu *et al*., 2023). By comparing the epigenetic landscapes of oysters from environments with differing levels of variability, researchers could investigate whether oysters exposed to highly fluctuating conditions exhibit distinct DNA methylation patterns, histone modifications, or small RNA-mediated regulation compared to their counterparts from more stable habitats. Experimental approaches could include assessment in common conditions, reciprocal transplant studies, or controlled laboratory exposures. This phenomenon could be evaluated further by isolating environmental effects from genetic background by exposing individuals from different populations to the same conditions and measuring epigenetic responses.

To date, there are limited studies demonstrating epigenetic modifications are more relevant in variable environments, as one might determine based on the suggested study above. However, there is evidence for constitutive upregulation where epigenetic factors have been attributed. A study in *Crassostrea virginca* notes that low pH changed DNA methylation and increased gene expression variability (i.e., transcriptional noise) in males while decreasing it in females (Venkataraman *et al*., 2024). More targeted studies are essential in this area.

#### 3.3.3 Mechanistic Knowledge Gap: What granularity of environmental change can organisms sense?

A corollary to our above hypothesis is that organisms should evolve epigenetically mediated responses to environmental changes that (1) recur frequently enough that the benefits of an epigenetic response outweigh their costs (and therefore the epigenetic response is evolutionarily maintained), and (2) are best addressed through reversible modification of gene expression. For example, experiments and models exploring the phenomenon of bet-hedging strategies (phenotypic variation that buffers an organism’s fitness in variable environments, (Reed *et al*., 2010)) demonstrate that these strategies should only evolve when the timescales of environmental variation are aligned with the timescales of an organism’s life history (Beaumont *et al*., 2009).

However, while a growing body of literature demonstrates a capacity for epigenetic remodeling in response to environmental change (Eirin-Lopez & Putnam, 2019), we still do not know the granularity of environmental change that organisms can sense (Dellaert & Putnam, 2023). Given evidence for both within-generation epigenetic modification (Dang *et al*., 2023; Gavery & Roberts, 2010; Putnam, 2021; Putnam *et al*., 2016, 2022; Rodriguez-Casariego *et al*., 2022) and trans-generational epigenetic inheritance (Liew *et al*., 2020; Putnam, 2021; Rondon *et al*., 2017; Strader *et al*., 2019; Wang *et al*., 2023), we posit environmental changes that occur once every two or fewer generations, but persist for at least one reproductive cycle, are those most likely to produce an evolved epigenetic response. However, these environmental changes should be reversible (Burggren, 2015), thus selecting for transient epigenetic modifications of gene expression, as opposed to permanent, directional evolutionary change.

By better understanding the frequency, speed, and environmental sensitivity of epigenetic modification experimentally and observationally, we can make hypotheses about the types of environmental variation that lineages experienced over evolutionary time.

## 4 Future Directions in Environmental Epigenetics

Our work identifies critical gaps in our ability to connect existing empirical data to theoretical predictions about how epigenetic modifications allow organisms to track their environments. Although affordable new sequencing technologies allow us to rapidly generate large amounts of data, we need to collect these data in the context of well-designed experiments if we are to successfully identify mechanisms. For example, as we detail above, to determine if and how organisms use information from their environments to preferentially add and remove epigenetic markers, experiments would ideally track the impacts of an environmental challenge (e.g., increases in temperature or decreases in pH) on organisms at physiological, cell biological, and epigenetic scales (Figure 4A). As more details are known about the timescale of epigenetic change, experiments can be designed more efficiently. By repeating these experiments in different populations and taxa, the field can work towards a unifying theory of epigenetic-environment linkages in marine invertebrates.

As empirical studies improve our mechanistic understanding, new mathematical models can be developed to incorporate these mechanisms. Generalized, phenomenological models like the one presented here can be used to unify findings into a broader framework, such as by identifying circumstances under which epigenetic markers might promote or inhibit gene expression. And tactical models tuned for specific systems can be used to understand the extent to which epigenetic modifications may buffer populations against directional environmental change. Of course, for these models to be accurate, rates and effect sizes of epigenetic changes must be well-constrained.

Overall, the field of environmental epigenetics is technologically and conceptually poised for major advances. Continued integration of mathematical frameworks with empirical data will help guide the field towards conceptual synthesis.

## Acknowledgments

The authors acknowledge support from the National Science Foundation Understanding the Rules of Life: Epigenetics program (EF-1921356 to HVM and RMN, EF-1921465 to HMP, EF-1921425 to RC, EF-1921149 to SBR, and EF-1921402 to JME-L). We are grateful for thoughtful feedback on early versions of the model and manuscript from Drs. Alexandra Brown, Ferdinand Pfab, and A. Raine Detmer.

## Supplementary Material

**Figure S1:**
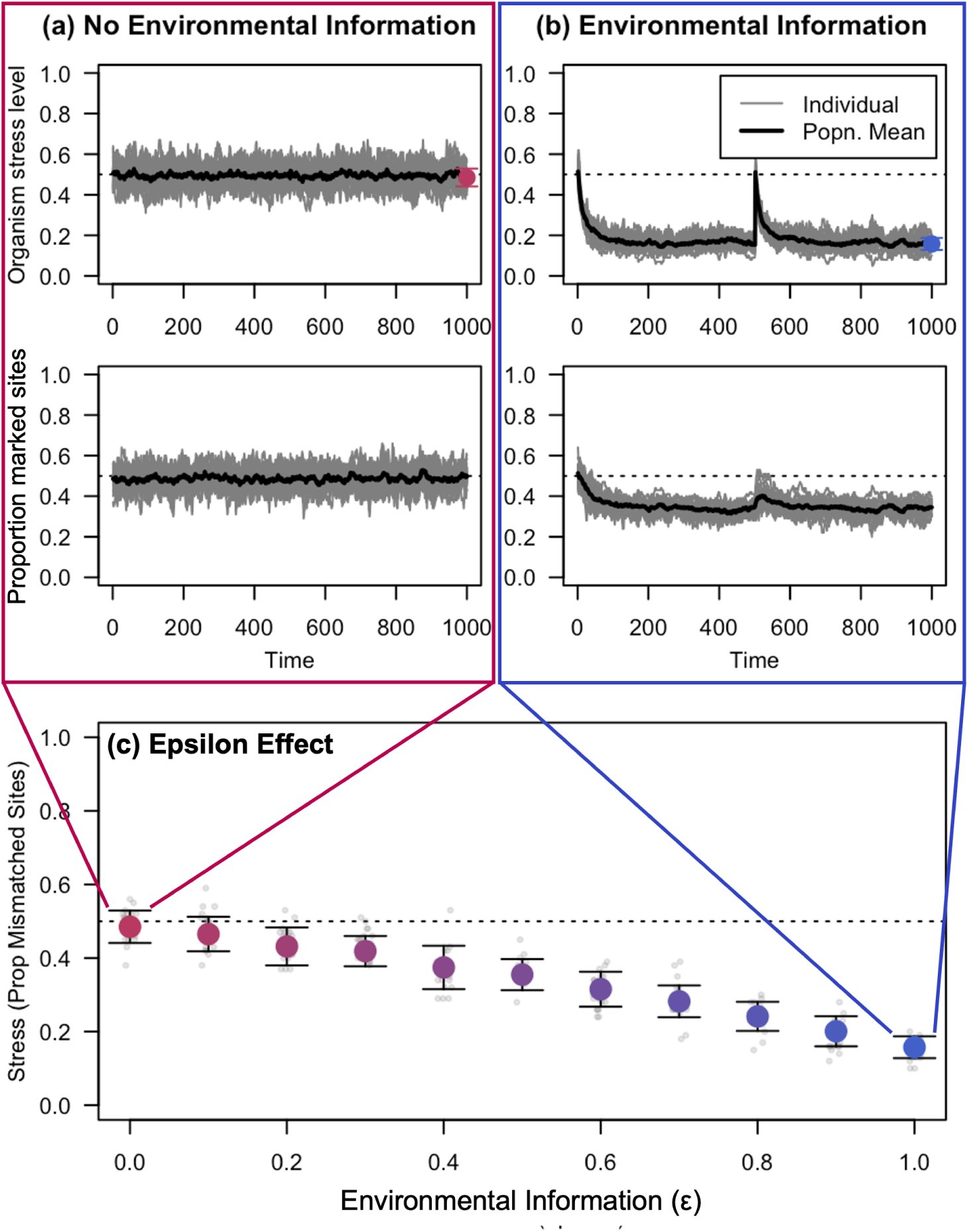
Effects of environmental information on organismal stress when marker addition is preferred. As in Figure 3: Top: Timeseries showing a group of 20 organisms failing to track (left) or tracking (right) the environment over time. Bottom: The larger the difference between preferential and baseline marker addition tendencies, the better the population’s performance. Simulations are for 20 indiviuals with a baseline marker removal tendency of 0.2 tracking an environment with a mean epigenetic marker abundance of 0.5. Bars show standard deviations around the mean.

**Figure S2:**
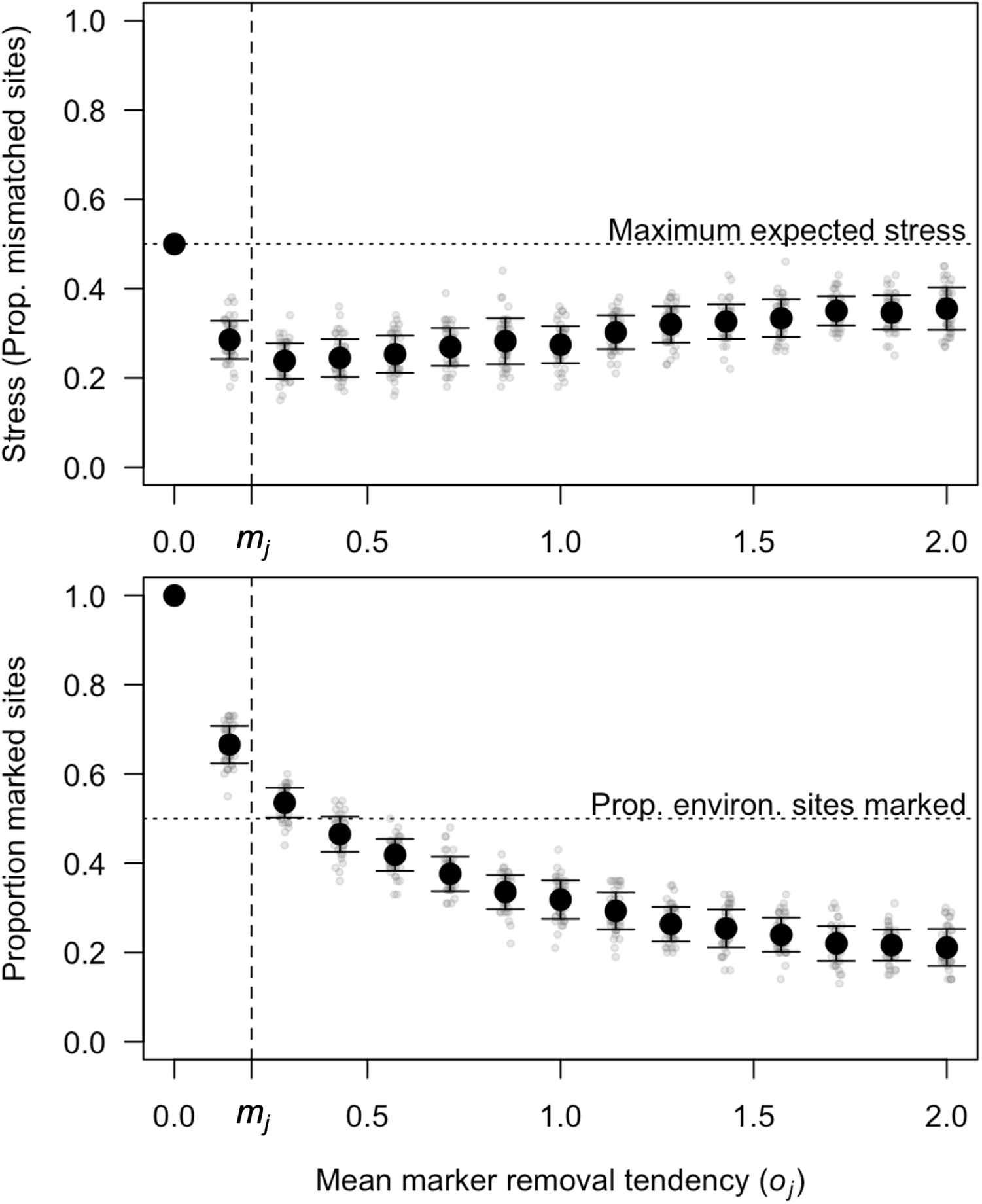
Dependence of stress and proportion sites with markers on marker removal tendency. Stress is minimized when mean marker removal tendency (o*_j_*) is similar to marker addition tendency (m*_j_*), resulting in the proportion of epigenetic markers on organism genomes slightly exceeding that required to match the environment (0.5). Simulations are for 20 organisms with a baseline marker addition tendency of 0.2 (vertical dashed line). Bars show standard deviation around the mean. The null expectation for stress and the proportion of environmental sites marked (0.5, dotted horizontal line) are also shown for reference.

**Figure S3:**
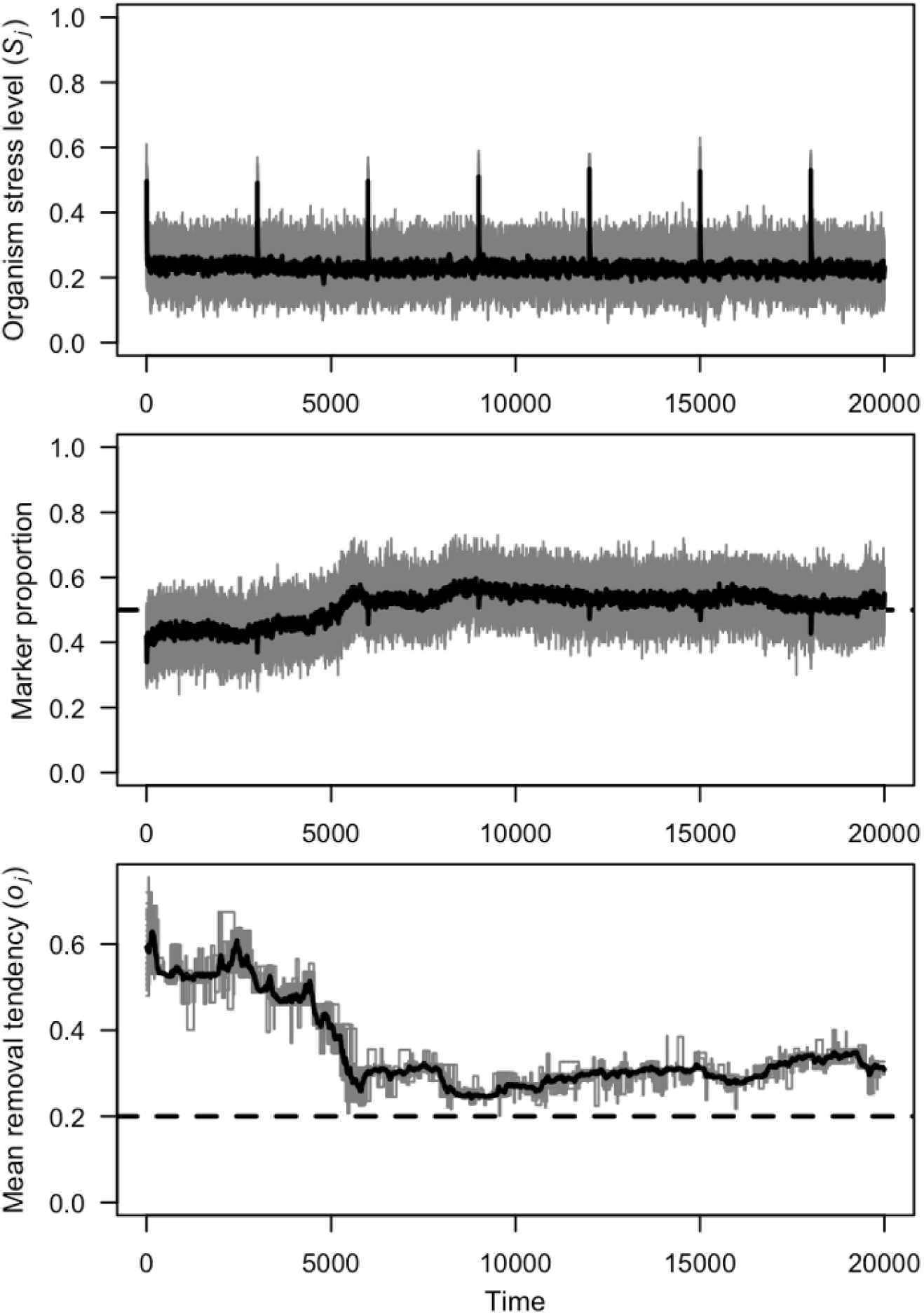
Trajectory of a population with a decreasing marker removal tendency over evolutionary time. The population is subjected to periodic environmental perturbations, which it recovers from, with decreasing stress as beneficial mutations accumulate. These mutations act to reduce the marker removal tendency and thus increase the overall genomic level of epigenetic markers to closer to the environmental mean of 0.5.

**Figure S4:**
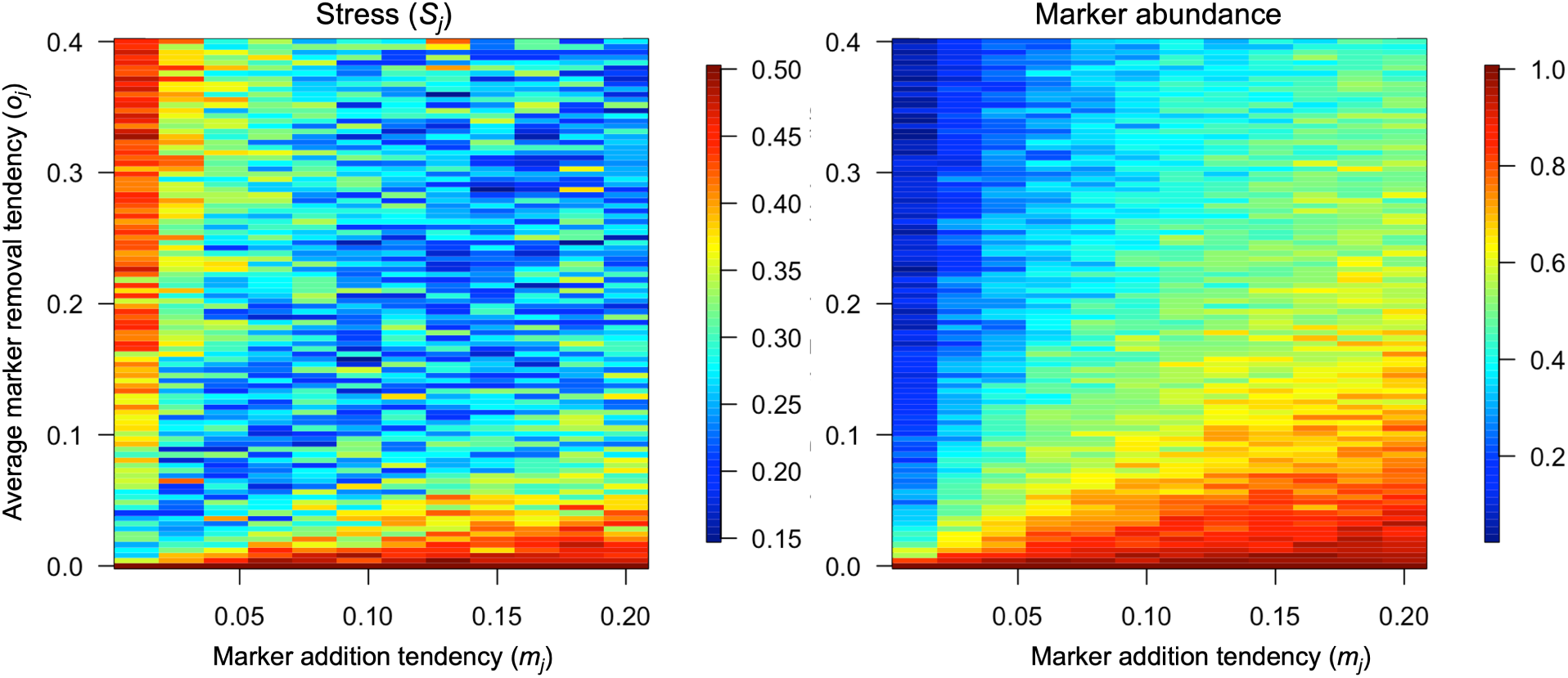
Effects of mismatched epigenetic tendencies on organism performance. If marker removal tendencies are much higher than marker addition tendencies (upper left of both panels), stress is high (left panel) because the organism’s number of epigenetic markers is low compared to the environment (right panel). In contrast, relatively low marker removal tendencies lead to high amounts of epigenetic markers on the genome and also high stress (lower right of panels).

**Figure S5:**
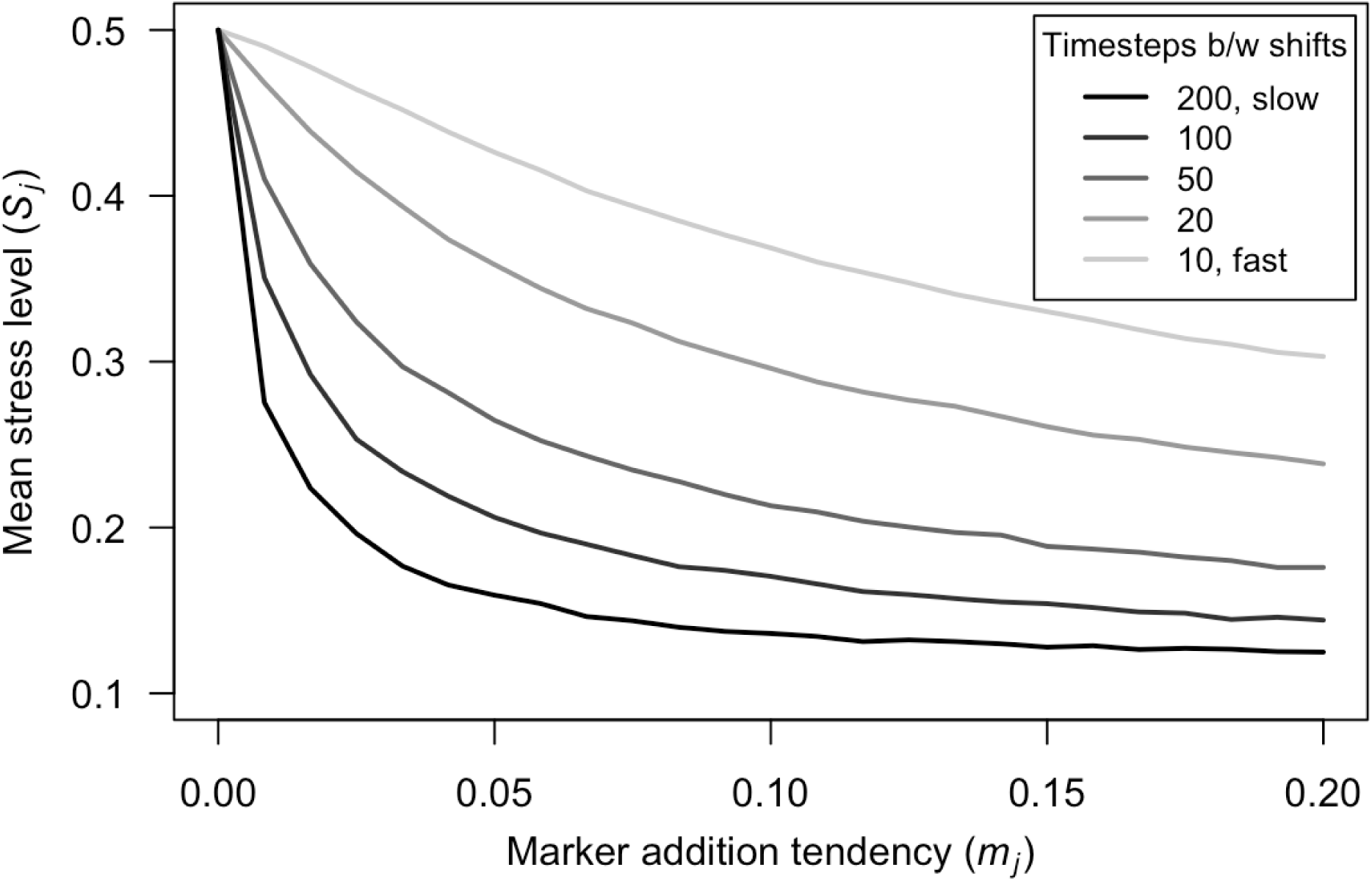
Relationship between environmental variability, organism traits, and experienced epigenetic stress. Here, we assume the environment is 50% marked, and that the marker addition and removal tendencies are optimally coupled following Figure 5b. As epigenetic modification tendencies increase, the number of modifications made per timestep increases. Given that some of the marker removal is preferential (ɛ = 0.8), greater tendencies lead to more effective tracking of the environment and lower stress levels. However, the effectiveness of environmental tracking also depends upon the environment: When the environment changes infrequently (e.g., once every 200 timesteps; black line), comparatively low epigenetic modification tendencies can nontheless reduce organismal stress levels. But more frequent environmental changes (e.g., every 10 timesteps; lightest gray line) result in high stress levels even when epigenetic modification tendencies are high. (Here, the magnitude of environmental change is 15 positions.)

